# A pan-eukaryotic analysis of mitochondrial ribosomes uncovers major evolutionary transitions

**DOI:** 10.64898/2026.10.06.756942

**Authors:** Tomáš Pánek, Jan Říha, Pia Corre, Eliška Zadrobílková, Harshini R. Vummadi, Arsh Suri, Prashant Chauhan, Jonathan E. Wong, Kira Senk, Anton S. Petrov, Victor Tobiasson, Ondřej Gahura

## Abstract

Mitochondrial ribosomes (mitoribosomes) have diverged extensively both from their bacterial ancestors and across eukaryotes, and while many general trends have been identified, fundamental features of their evolution remain unclear. We analyzed sequences from 85 eukaryotes to chart protein composition across the evolution of mitochondrial translation. We both reconstruct the LECA mitoribosomal proteome as containing nearly the full bacterial protein complement and at least 23 proteins recruited after α-proteobacterial endosymbiosis and describe the subsequent extensive but lineage-specific remodeling post LECA. We further uncover previously unrecognized homologies among several mitoribosomal proteins and unify and update the mitoribosomal nomenclature. Our data also support protein retargeting between organellar ribosomes, documenting remarkable intercompartmental structural compatibility. Together, these findings highlight the exceptional evolutionary plasticity of mitoribosomes and provide a comprehensive reference framework for future analysis.

## INTRODUCTION

Protein synthesis is catalyzed by large ribonucleoprotein complexes known as ribosomes (Fig. 1A). In eukaryotic cells, ribosomes are localized not only in the cytosol, but also in mitochondria and plastids, endosymbiotic organelles which retain their own genomes and gene expression machineries. Mitochondrial genomes encode the ribosomal RNAs (rRNAs) and a limited subset of mitochondrial ribosomal proteins while most mitoribosomal proteins are encoded in the nuclear genome(1). Given the essential role of mitochondria in cellular metabolism, understanding the evolution of the organellar translation machinery across eukaryotes is not only important from an evolutionary perspective but can also inform targeted therapies against pathogens with divergent mitoribosomal features. Furthermore, mitoribosomes, and their role in organellar protein synthesis, represents a valuable model for studying the evolutionary transition from a bacterial endosymbiont to a fully integrated organelle.

**Figure 1:**
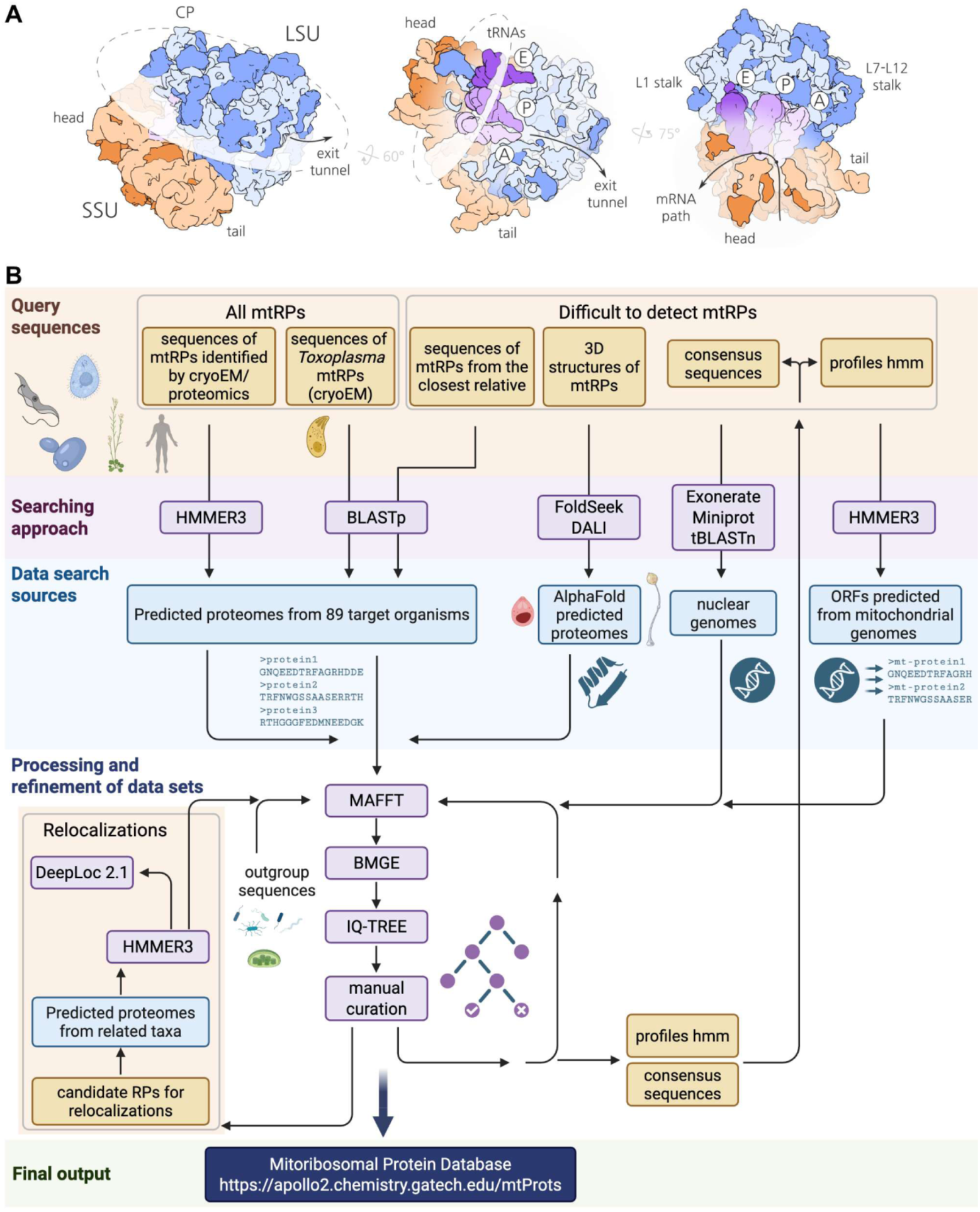
A scheme of ribosomes and workflow of search of mitoribosomal proteins. **(A)** A general scheme of ribosomes and their subunits viewed from intersubunit sides with functional and morphological features. The large and small subunits are shown in blue and orange, respectively. Proteins are in dark shades, RNA in pale. tRNA molecules are shown in shades of purple. CP; central protuberance. (B) Overview of the workflow for ortholog detection, filtering, and validation employed in this study. The panel (B) was Created in BioRender. Gahura, O. (2026) https://BioRender.com/cjuvd2k

The uniformity of ribosomes in extant bacteria implies that mitoribosomes evolved from an ancestral state highly similar to the bacterial form (2). Early biochemical and subsequent genomic and proteomic studies (reviewed in (3–5)) indicated that mitoribosomes have diverged markedly from their bacterial counterparts. Subsequent structural characterization by cryoelectron microscopy (cryoEM) revealed extensive divergence in their protein composition and overall architecture (6–18). These structural studies provide the most reliable means of defining the protein complement of individual mitoribosomes and were crucial in revealing previously unrecognized shared ancestries (homologies) between proteins with high sequence divergence. However, cryoEM is not a feasible approach for most eukaryotic microbes, as only a few can be cultured at sufficient scale (19).

Mitoribosomal proteins can be classified into three groups based on their origin: (1) proteins shared with bacterial ribosomes, (2) proteins acquired after the endosymbiotic event and before the emergence of the last eukaryotic common ancestor (LECA), and (3) lineage-specific proteins acquired later in the evolution of eukaryotes. While some proteins of bacterial origin remain encoded in mitochondrial genomes, proteins from the latter two groups are encoded in nuclear genomes. Lineage specific proteins have been recruited from proteins found in the mitochondrial matrix, such as enzymes or RNA binding proteins, or emerged as paralogs of existing ribosomal proteins, neofunctionalized in the context of the ribosome as novel structural components (16,18,20).

Here, using a combination of bioinformatic approaches with extensive manual curation, we integrate recently available high-resolution structural information with genomic data from representatives of all main eukaryotic lineages to systematically map mitoribosome diversity across eukaryotes. We infer the mitoribosomal protein composition in nearly all major eukaryotic lineages, reconstruct the mitoribosome of the LECA, and define the principal trends and constraints that shaped the evolution of mitoribosomal architecture.

## MATERIALS AND METHODS

### Input data resources

The list of target organisms contained 64 species with high-quality genomes and, to fill gaps in the eukaryotic diversity, additional 21 organisms for which only transcriptomes are available. In total, we searched predicted proteomes from 85 species, covering the entire eukaryotic tree of life. From these, seven species do not contain mitochondrial genomes and mitoribosomes, and they serve as negative controls. Further, we searched for ribosomal proteins encoded in mitochondrial genomes of additional 36 species. For summary of target organisms and sequence data sources, see Supplementary Table S1. To query predicted proteomes of target species and to build the initial protein datasets, we collected sequences of mitoribosomal proteins from humans (10,21), *Saccharomyces cerevisiae* (9,22), *Arabidopsis thaliana* (14), *Tetrahymena thermophila* (8) and *Trypanosoma brucei* (18), whose mitoribosomes have been characterized structurally, and from three species, whose complements of mitoribosomal proteins were previously reported based on proteomic (*Andalucia godoyi* (20)) or biochemical (*Acanthamoeba castelanii* (23) and *Euglena gracilis* (24)) studies.

### General search workflow, filtering and validation

We used the query sequences to search the sequence data from target organisms by profile HMM based strategy, complemented by structure similarity-based approach in selected cases, and performed extensive manual filtering and curation of the data using phylogenetic analyses, and visual inspection of multiple sequence alignments or predicted structures of selected candidates to remove non-mitoribosomal proteins and contaminations from other species. We performed these steps iteratively to further improve profile HMMs (see Figure 1B for an overview of the workflow).

Sensitivity and specificity of our search were validated by comparing our results with published cryoEM studies of mitoribosomes from *Chlamydomonas reinhardtii* (13) and *Toxoplasma gondii* (16,17). Our search detected only proteins previously found in the structures or those that localize in regions with high flexibility or low resolution of the respective cryoEM map. This validates the high specificity of our approach. However, we did not find several proteins identified in mitoribosome structures, implying that the search sensitivity might be insufficient for generally divergent proteins if query sequences from a related organism are not available. Therefore, we classified proteins that can remain undetected by the original approach due to their short length, high divergence or existence of non-ribosomal paralogs (marked as “difficult to detect” in Fig. 2) and performed additional searches. First, structures of uL10m, bS21m, mS22, mS26, mS37 and mS38 from *A. thaliana*(14) were used to query structure databases. We employed DALI server (25) and FoldSeek (26) to search the AlphaFold (27) database of predicted structures of complete proteomes of two target organisms, *Plasmodium falciparum* and *Dictyostelium discoideum*, for the presence of proteins with similar structure. This approach resulted in identification of uL10m in *D. discoideum.* We also searched original nucleotide assemblies of nuclear and mitochondrial genomes to exclude the possibility of missing mitoribosomal proteins due to incomplete genome annotations (for details, see Figure 1B and below).

**Figure 2.**
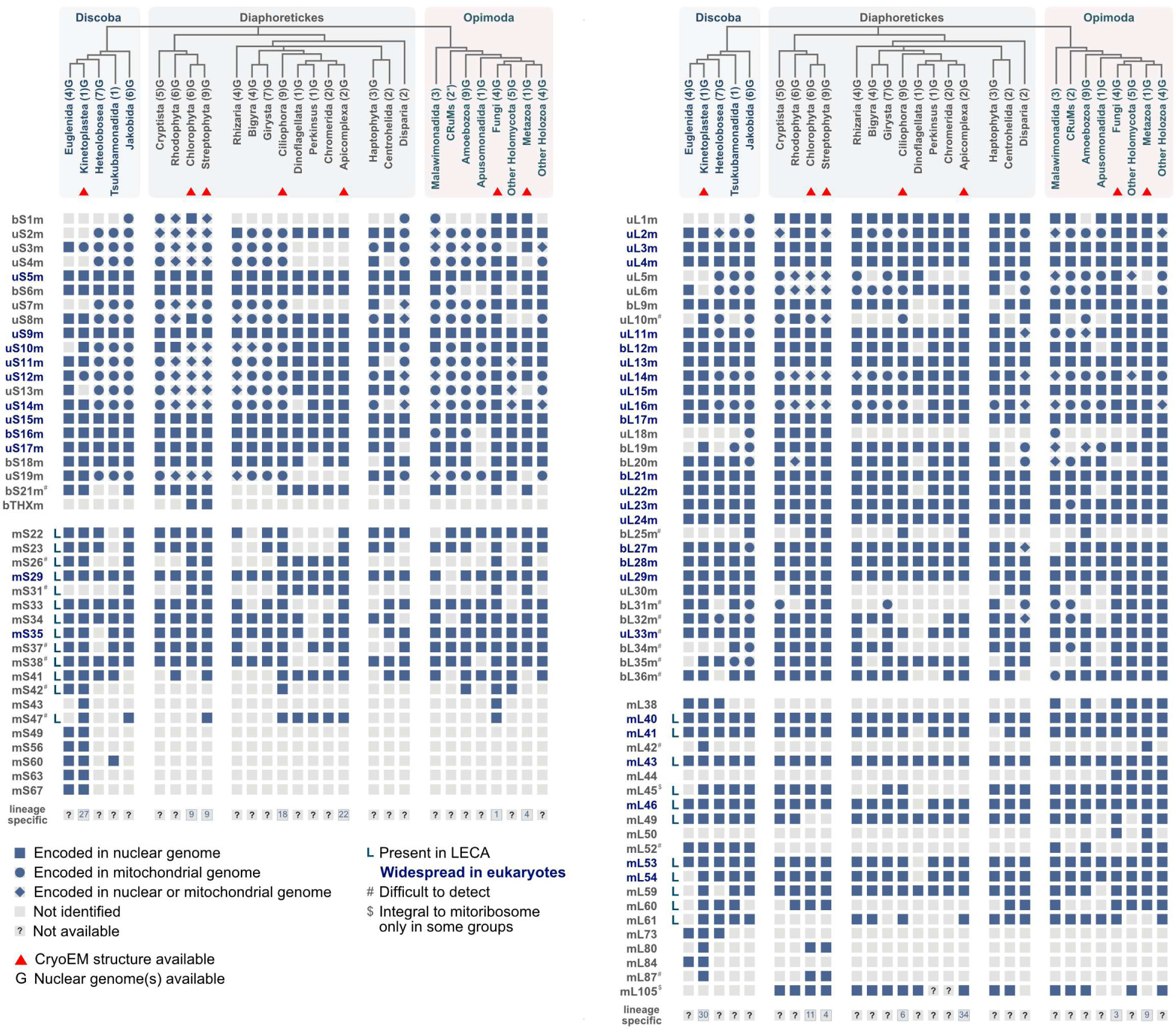
Distribution of mitoribosomal proteins. Numbers in parentheses indicate numbers of organisms in individual groups. Schematic phylogenetic tree is based on (59). Distribution of mL105 is based on (98). Presence of bS6m in CRuMs is based on (69). We classify short mitochondrial proteins that can be overlooked during gene prediction, and proteins difficult to distinguish from non-mitoribosomal homologs, even with comprehensive phylogenetic analyses, as “difficult to detect”.

### Analysis of difficult-to-detect and relocalized proteins

We performed additional analyses of difficult-to-detect mitoribosomal proteins (Fig. 2) using iterative search in nuclear and mitochondrial genomes and transcriptomes of 54 selected target organisms (see Supplementary Table S2) with profile Hidden Markov Models (HMMs) of selected mitoribosomal proteins, followed by phylogenetic analyses with IQ-TREE v2.3.5 (28) and targeting predictions with DeepLoc 2.1 (29). Profile HMMs(30) of mitochondrial ribosomal proteins were constructed from the datasets generated in this study. Multiple sequence alignments for profile HMMs construction were computed using MAFFT L-INS-I (31) (for detailed settings see below). Putative open reading frames (ORFs) in mitochondrial genomes were identified de novo and translated by TransDecoder (https://github.com/sghignone/TransDecoder), using the genetic code specified in the original NCBI annotation and a minimum ORF length of 40 amino acids. Putative ORFs were subsequently used for homolog search using profile HMMs of mitochondrial ribosomal proteins that were assigned as difficult-to-detect, followed by iterative phylogenetic analysis with IQ-TREE v2.3.5. Nuclear genome assemblies were searched using exonerate (32), settings: model protein2genome, score variable upon length of protein from score 70 for short proteins to 125 for longer proteins), miniprot (33), settings: minimal ORF length 20, gap penalty set to 7), and tBLASTn (34) (settings: E-value threshold variable upon protein length from 1e-7 to 1e-11). For miniprot setting, and for retrieving sequences from exonerate and tBLASTn, we considered nuclear genetic codes described in Supplementary Table S1. Profile HMM consensus sequences generated from these profiles using *hmmemit* (plurality-rule consensus sequence obtained with *-c* option) (30) were used as queries whenever possible. For highly divergent proteins, sequences from the closest available taxonomic relative were used instead. Candidate protein sequences identified in nucleotide assemblies were filtered by phylogenetic analyses as described above. Curated sequences were subsequently incorporated into the ribosomal protein database.

For analysis of protein relocalizations, i.e. events, when a ribosomal protein replaced its counterpart in the ribosome of other cellular compartment, we iteratively enriched our set of target organisms by relative selected species and, where applicable, by organisms where relocalizations were suggested by previous analyses. This approach allowed us to estimate, when in evolution the relocalization events of individual proteins occurred. We searched nuclear, mitochondrial and plastid genomes of these organisms with mitochondrial, plastidial and cytosolic profile HMMs using HMMER3 (30), or using BLASTp (35) with a query sequence from closely related organisms. The profile HMMs of non-mitochondrial ribosomal proteins were constructed from annotated sequences in plastid genomes on NCBI RefSeq and plastidial and cytosolic ribosomal proteins identified by BLASTp search of EukProt v3 (36) with *A. thaliana* sequences as a query. For several genomes relevant for the analysis without available gene annotations, we used exonerate search (model protein2genome, threshold score 80) to identify candidate ORFs corresponding to ribosomal proteins of interest. Targeting predictions for all nuclear encoded proteins was carried out with DeepLoc 2.1 (29). All organisms and genomic resources are summarized in Supplementary Table S1.

### Phylogenetic analyses

Protein sequences were aligned using MAFFT v7.453 (31) with the following settings: L-INS-i algorithm, gap opening penalty 1.3, leave gappy regions, and the BLOSUM30 scoring matrix. The alignments were trimmed either manually or using BMGE v1.12 (-b 2 -g 0.8)(37). Phylogenetic analyses were performed with IQ-TREE v2.3.5 (28) using the LG4X model and 1,000 ultrafast bootstraps with bnni correction (the model was selected arbitrarily as a reasonably complex model, given the size of the matrices fed into the phylogenetic analyses). All protein sequences, untrimmed and trimmed alignments, and raw phylogenetic trees are deposited on Figshare.

For final phylogenetic trees documenting protein relocalizations, duplicated sequences were filtered, the dataset was enriched by homologs from close relatives, aligned using MAFFT 7.526 (31) with settings as described above and manually trimmed. Fifteen independent maximum-likelihood searches were performed using IQ-TREE v2.3.5 (28) under LG+C60+G4 (settings with *-mwopt* option enabled and *-nstop* increased to 150). We employed the complex and computationally expensive profile mixture model C60 (38) for the final tree inference, because this model accounts for site heterogeneity, reducing the risk of long-branch attraction artifacts. Recent studies have also shown that C60 can effectively handle short alignments without over-parameterization (39). The topology with the highest log-likelihood was selected as the best scoring tree. Branch support was assessed by 5000 ultrafast bootstraps with bnni correction and SH-alrt test with 5000 replicates. Predictions of subcellular localizations to Mitochondria or Plastid assessed by DeepLoc 2.1 (29) were mapped onto phylogenetic trees with their prediction scores by a custom python script and trees were visualized in FigTree 1.4.4 (https://tree.bio.ed.ac.uk/software/figtree/).

In this study, we utilized OpenAI ChatGPT (versions 5 to 5.5, August 2025–May 2026) to assist in the generation and review of scripts for data analysis. All AI-generated content was critically evaluated, edited, and validated by the authors to ensure accuracy and integrity in the published results.

### Database

MitoRPdb (https://apollo2.chemistry.gatech.edu/mtProts) is a web-based tool that aims to share the information about mitoribosomal proteins across 85 eukaryotic species included in this study. Each entry contains sequence, genome location, assembly ID, alternative names and available 3D structures (along with the corresponding PDB and chain IDs). The tool allows filtering records using several queries and downloading individual sequences and multiple sequence alignment for user-defined selection of proteins and organisms. MitoRPdb was designed using a client–server architecture; the web server is hosted at Georgia Institute of Technology and served by Apache/RHEL8. The backend is powered by Python 3.8 and uses Django for API queries and data processing. The front-end user interface was developed using HTML5/JavaScript, leveraging the React and Vue frameworks. The data is stored in a MySQL database. MitoRPdb also relies on information provided by several external APIs and databases.

### Modeling of mitoribosome in LECA

Proteins with bacterial origin are represented by structures of respective *E. coli* homologs (PDB ID 7K00 (40)). In cases of three mtSSU and ten mtLSU proteins, extensions present in LECA were deduced from MSAs of extant bacterial and mitochondrial ribosomal proteins. The structures from *A. thaliana* were used to depict the extensions in LECA as well as the cores of mitochondria-specific proteins inferred from MSAs to be present in LECA. The structures from *A. thaliana* were chosen because out of all structurally characterized mitoribosomes, the one from this organism most closely resembles bacterial ribosomes (14). To visualize the 3D structure of LECA mitochondrial rRNA, we used *E. coli* rRNA structure, because structure of a ribosome from α-proteobacteria, the closest extant relatives of mitochondria, is not available. To compensate differences between rRNA of *E. coli* and mitochondria, we removed LSU helices H10, H58-59 and H98, and the terminal hairpin of SSU helix h17, which are missing in all structurally characterized extant mitoribosomes and the latter three also in α-proteobacteria. Further, we do not depict LSU helices H55-H57, whose ancestral state cannot be unambiguously inferred from the structures of mitoribosomes due to combination of poor sequence conservancy, local structural shifts and frequent lineage-specific expansions. 5S rRNA highly similar to that of extant bacteria was present in the mitoribosome of LECA (41) and is also included in our reconstruction.

## RESULTS AND DISCUSSION

### *In Silico* framework for detecting mitoribosomal proteins

To survey the distribution of mitoribosomal proteins across eukaryotes, we queried genomic and transcriptomic data from 85 strategically sampled organisms with the protein sequences present in at least two of the nine species with structurally or proteomically characterized mitoribosomes. We employed an iterative, phylogeny-aware ortholog detection pipeline supplemented with rigorous manual curation and reannotation of genomic data (see Fig. 1B and Methods for details). To minimize the effect of incomplete genomic data, each major eukaryotic group was represented by several high-quality genomes or transcriptomes (see Supplementary Table S1). Our approach outperforms previous *in silico* analyses and corrects misannotations included in previous studies focused on individual organisms (23,24,42), exemplified by our ability to assign identity to previously uninterpreted regions in cryoEM maps (bL34m and mS31 in *Arabidopsis thaliana*, and five proteins in *Chlamydomonas reinhardtii*; Supplementary Fig. S1, Supplementary Table S2).

Using this framework we also extend previous sequence, structure, and interaction network analyses to indicate several unnoticed homologies among mitoribosomal proteins (20,43) (Table 1 and Supplementary Table S3). We note proteins previously termed mL60 in *S. cerevisiae* and *A. thaliana*, and mL63 in humans and trypanosomes are orthologs, and that mL88 in *T. brucei* is an ortholog of mL59 (Supplementary Fig. S2A). Furthermore, the human and *T. brucei* mL42 are likely orthologous to *S. cerevisiae* mL58 (Supplementary Note 1; Supplementary Fig. S2B). Several mitoribosomal proteins originated in gene duplication events, as exemplified by human mL66 and mS40, which are paralogs of *bS18m* gene (previously termed *MRPS18A*, *MRPS18B*, and *MRPS18C*, respectively (44)). We document two cases of previously unrecognized paralogy in euglenozoan mitoribosomes: mL84 is a paralog of bL9m, and proteins mS50, mS58, and mS60 are all distant paralogs (Supplementary Fig. S3). Finally, we observe that bL12m, the protein oligomer forming part of the L12 stalk, has been duplicated at least once in four independent lineages (myzozoans, diatoms, spermatophytes, and kinetoplastids; Supplementary Fig. S4) possibly producing a generally heteroligomeric arrangement, as supported by the presence of both paralogs in mitoribosomes containing samples from *T. brucei* (18,45). To unify naming of all hidden orthologs and paralogs revealed in this study or elsewhere (20,43) and to take account of the prevalent trend of protein fragmentation, we propose an update and extension to the established mitoribosomal protein nomenclature (46,47) (Supplementary Note 2 and Supplementary Table S3). To further aid in the analysis of mitoribosomes we provide a reference script compatible with ChimeraX with all proteins labeled in accordance with this study (Supplementary Data 1).

**Table 1.** List of paralogs within mitoribosomal proteins.

| Paralogs | References |
| --- | --- |
| bS18m, mS40, mL66 | (44) |
| mL37, mL65 | (6) |
| uS10m, mL48 | (64) |
| mS22, mL42/mL58 | (20) |
| mS25, mL43, mL53, mL61, mL159 | (16,20) |
| uL24m, mL134 | (16) |
| mL132, mL135 | (16) |
| mS135, mS136 | (16) |
| bL9m, mL84 | this study |
| mL80, mL164 | this study |
| mS50, mS58, mS60 | this study |
| mS35, mL62, (mitochondrial) release factors | (93); this study |

### Mitoribosomal proteins with (near-)universal distribution

A summary of identified mitoribosomal proteins across all examined organisms is provided in Supplementary Table S2. All identified protein sequences with metadata are available in a newly developed web database MitoRPdb (https://apollo2.chemistry.gatech.edu/mtProts; see Methods for details). To reveal global trends in eukaryotic evolution, we pooled data from individual eukaryotic organisms into higher-order phylogenetic groups (Fig. 2). We classified 35 mitoribosomal proteins present in all reported structures and absent in no more than two of these groups as “widespread” (Fig. 2). These proteins are ubiquitous constituents of the ribosome filling universal functional roles, and unlikely to be missing even in the most divergent mitoribosomes.

Out of the widespread mitoribosomal proteins, 27 are of bacterial origin (Fig. 2). This group includes all proteins that line the polypeptide tunnel and its exit in bacteria (uL4, uL22, uL23, uL24 and uL29), components of the L12 stalk interacting with elongation factors (bL12 and uL11) and proteins crucial for fidelity of decoding (uS5, uS11, uS12 and uS17) (48). In addition, essentially all proteins that bind to rRNA early in the assembly process of bacterial ribosomes and mitoribosomes, as inferred from multiple lines of evidence(49–53), are also ubiquitous and suggests that early biogenesis of mitoribosomes is conserved.

Out of 8 widespread mitoribosome-specific proteins (Fig. 2), three (mS29, mL40, and mL46) universally contribute to contacts between the small mitoribosomal subunit (mtSSU) head and the central protuberance of the large subunit (mtLSU) (10,15,54). mL40 also wraps around the central protuberance, irrespective of its composition, possibly providing reinforcement or aiding assembly. Proteins mL41 and mL43, while structurally disparate, colocalize with, and carry charge profiles compatible to early mt-rRNA reductions, and were therefore likely recruited early in the evolution of mitoribosomes (Supplementary Fig. S5). mS22 bridges two ancestral proteins of bacterial origin, bS16m and uS5m, via structurally conserved contacts; however, it remains undetected in several major groups, likely due to its remarkable structural variability (Supplementary Fig. S5A). Protein mL59 connects the base of the L1 stalk and the central protuberance, and extensions of mL59 have been shown involved in L1-stalk/tRNA interactions (54).

Two additional proteins, mS37 and mS38, play a central role in the maturation of the decoding center in mammalian mitoribosomes (55) and their interaction network is conserved in the trypanosomal mtSSU precursors (56,57), indicating an ancestral and broadly conserved function in mitoribosome biogenesis. Because these two proteins are present in all query organisms (including *C. reinhardtii*, where we identified them in this study), we consider them widespread despite difficulties in detection due to their short length. Likewise, three proteins (uL10m, bS21m, and mS26) present in all query organisms, were not detected in several target species likely due to the high degree of sequence divergence among lineages (Supplementary Data 2). However, we assume their widespread distribution, in line with their key roles in the ribosome. Specifically, bS21m co-constitutes mRNA channel, uL10m forms a part of the base of the L12 stalk and direct interactions with bL12m, responsible for recruiting translation factors to the ribosome.

Several other proteins, regardless of their origin and phylogenetic distribution, are generally difficult to detect. These proteins are marked in Fig. 2 and details are provided in Supplementary Note 1.

### Reconstruction of mitoribosomes in LECA

The reconstructed distribution of mitoribosomal proteins allowed us to infer the protein composition of mitoribosomes in the LECA. For the inference, we used a conservative approach based on the existence of three clearly monophyletic groups of eukaryotes (58,59) (Fig. 2). Proteins found in at least one organism in Discoba, Diaphoretickes, and Opimoda were inferred to be present in the LECA. Based on our analysis the LECA mitoribosome contained 54 proteins of bacterial origin, i.e. all proteins found in eubacteria except for bS20, and at least 23 additional mitochondria-specific ribosomal proteins. If the eukaryotic root is placed between Opimoda and the rest of eukaryotes (Diphoda) as indicated by some analyses (60,61), the number of mitochondria-specific proteins in LECA would increase to 26 (additional proteins are mL38, mL42, and mL52). Regardless of rooting, our analysis significantly expands the LECA mitoribosomal proteome compared to earlier studies which predate published cryoEM structures (20,62–64).

Our reconstruction documents that while the rRNA of the LECA mitoribosome was essentially α-proteobacterial-like, the protein content was substantially higher than in extant α-proteobacteria, attributed to both acquired proteins and ribosomal protein extensions (Fig. 3A). The acquisition of new proteins was not accompanied by loss of ancestral proteins, with the sole exception of bS20, routinely lost in diverse bacterial lineages and possibly absent from the α-proteobacterial ancestor (65,66). During eukaryogenesis, the compositional complexity of the mitoribosome increased more markedly than that of the cytosolic ribosome. Assuming that at the time of α-proteobacterial endogenization the protoeukaryotic cell possessed cytosolic ribosomes comparable in complexity to extant Asgard archaea, then during the first eukaryotic common ancestor (FECA) to LECA transition the cytosolic and mitochondrial ribosomes diverged in complexity, acquiring 11 and 23 proteins respectively. In addition, the nature of those acquired proteins differ, with primarily small globular folds of low structural complexity being added to the cytosolic ribosomes, compared to a wider diversity of protein folds on the ancestral mitoribosome.

**Figure 3.**
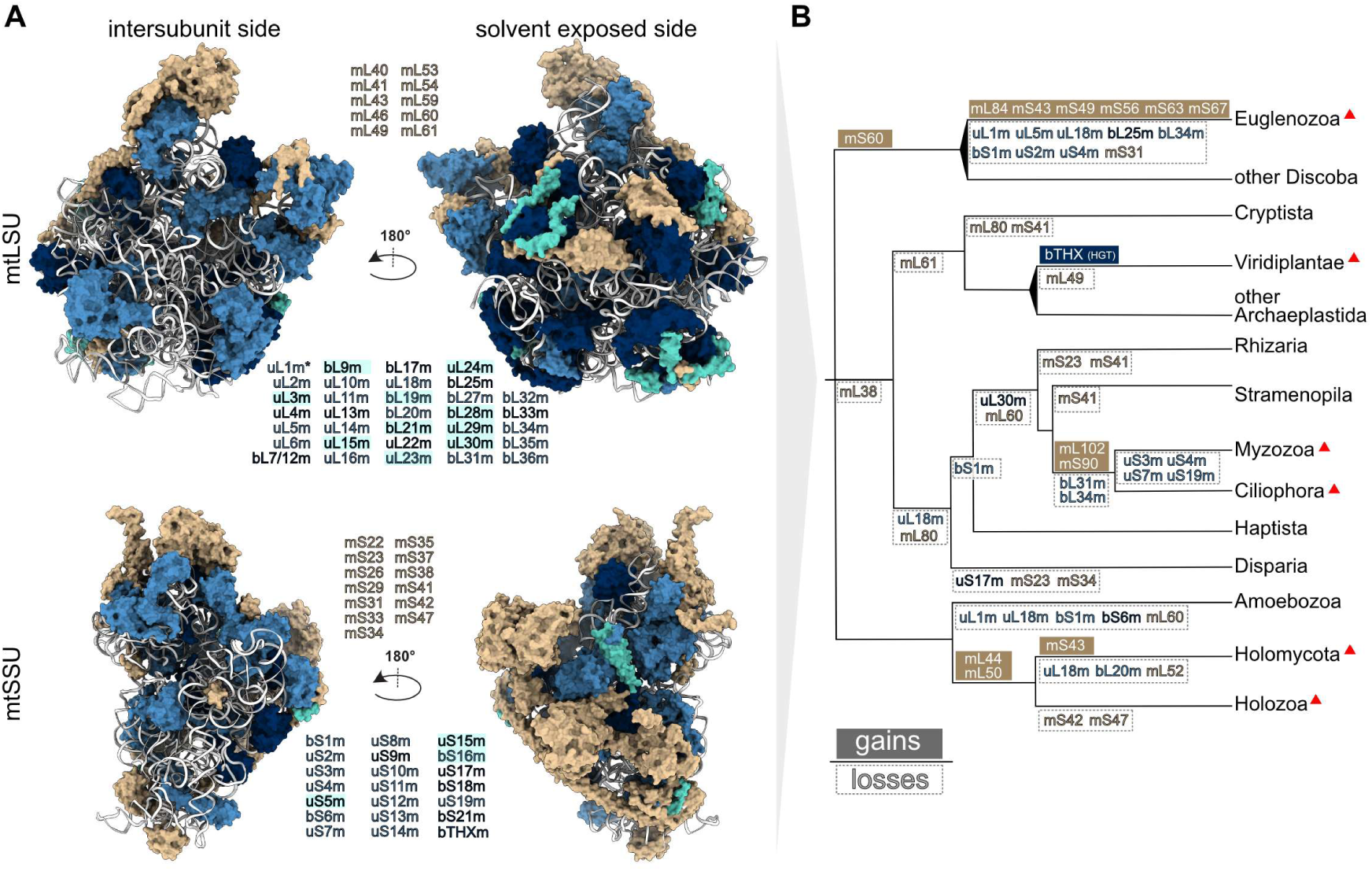
Mitoribosome in LECA and evolution of protein composition of mitoribosomes in eukaryotes. **(A)** Reconstruction of the mitoribosome in LECA. RNA is shown as white ribbon, proteins of bacterial origin encoded in mitochondrial and nuclear genome of LECA are shown in blue and navy, respectively, extensions of proteins of bacterial origin are cyan, and mitochondria-specific proteins are tan. **(B)** Gain and losses of mitoribosomal proteins mapped on the phylogenetic tree of eukaryotes as in Fig. 2. Gains and losses are shown above and below individual branches, respectively. Proteins are colored as in (A). Losses of proteins classified as “difficult to detect” (Fig. 2) are shown only for groups with available structure, which are indicated with a red triangle. Gains of proteins specific for organisms with cryoEM structures are not shown. bTHX was gained by horizontal gene transfer (HGT) from bacteria.

In summary, whereas the protein component of the mitoribosome already expanded in the pre-LECA period, significant changes in the mitoribosomal rRNA occurred after LECA, in parallel with diversification of protein content by lineage specific acquisition and losses

### Structural replacements or losses aid diversification of mitoribosomes

Post-LECA, mitoribosome evolution was marked by extensive gains and losses of protein components (Figs. 2&3B), collectively giving rise to lineage-specific features. While acquired proteins often originate from duplications of proteins found in the mitochondrial matrix, some proteins, i.e. mS93 (mTERF) in *T. thermophila* (8) and mS47 (hydroxyisobutyryl-coenzyme A hydrolase) in *S. cerevisiae*(9), have been entrenched without prior duplication. These proteins simultaneously act as ribosomal proteins while presumably retaining their original function. We observe that most losses occurred independently multiple times, exemplified by bS1m, uS4m or uL18m, and that losses of broadly conserved proteins (e.g. uL1m, uS8m or uS13m) are restricted to lineages with generally divergent mitoribosomes, such as Alveolata or Euglenozoa (67).

Changes in lineage-specific protein composition were preceded by convergent RNA reduction in several eukaryotic lineages, notably including Metazoa. Through a mechanism referred to as patching, it was proposed that functions of lost RNA elements were replaced by lineage-specific protein features (68), driving structural divergence (64). The parallel replacement of mtLSU rRNA region H53-59 by various proteinaceous structures in human, fungal, and trypanosomal mitochondria (Supplementary Fig. S6A) were given as examples of this proposed general trend. However, many seemingly critical and well conserved RNA elements were not positionally replaced by proteins, as exemplified by the loss of helices H6 and H7 by the polypeptide exit tunnel in humans (21) and *Trypanosoma* (18) (Supplementary Fig. S6A), nor does protein recruitment specifically occur in regions undergoing rRNA reduction. As more structures are determined the lack of correlation between rRNA loss and protein acquisition becomes more apparent. For example, in *Polytomella* and *Chlamydomonas*, large rRNA reductions have occurred without associated protein recruitment(7,13), and in *Tetrahymena* and *Toxoplasma*, the vast majority of lineage-specific proteins is recruited in absence of any major rRNA losses (8,16). Finally, patching does not address how these features proliferate within a population as deleterious losses of rRNA are unlikely to fixate. Taken together, while extensive protein entrenchment has followed rRNA losses in all studied mitoribosomes, it is highly unlikely that this process is driven by adaptive evolution aimed at patching lost rRNA.

The evolutionary trajectories following protein losses are likewise diverse. Some losses are accompanied by positional replacement, e.g., bL20m is replaced by mL58 in Holomycota (Fungi and their unicellular relatives) or uS2m by mS59 in trypanosomes. Other lost proteins are replaced only in certain groups, while their positions remain unoccupied in others. A representative case is uS13m, which has been replaced by a similar structure constituted by mS49 and mS71 in trypanosomes, but its position is empty in the human mitoribosome (Supplementary Fig. S6B). Some proteins have been lost even though the regions of RNA they contact remain structurally unchanged. Conversely, other proteins are retained, although their original interacting rRNA interfaces are reduced or completely absent. Both situations can be illustrated by uL30m. In bacterial ribosomes, uL30 sits in a hollow formed by the largest internal loop of the LSU RNA domain II (single-stranded regions connecting helices H36-41&45) and forms no contacts with other proteins. This arrangement is retained in mitoribosomes from plants, fungi, and mammals. In trypanosomes, the RNA domain II loop is highly reduced, but the core of uL30m remains unchanged and is surrounded nearly exclusively by lineage-specific proteinaceous elements. In contrast, uL30m was lost in an ancestor of alveolates, even though both the structure and the sequence of all associated RNA elements are conserved (Supplementary Fig. S6C). Similarly, bL34m, a small protein enclosed in the mtLSU by RNA elements of domains I-III, is lost without compensation in mitoribosomes of *Tetrahymena*, although its binding pocket is largely conserved. bL34m is also missing in the model of the mitoribosome of *A. thaliana* (14) (PDB ID 6XYW). However, our search identified a candidate protein, which fits into unmodeled density in the respective cryoEM map (EMD 10654; Supplementary Fig. S1), aiding the completeness of the model. Taken together protein recruitment is a highly dynamic process which, while coinciding with both protein and rRNA losses, is unlikely driven by gain or restoration of function and appears to follow principles of a neutral complexity ratchet.

### Endosymbiotic transfer of genes encoding mitoribosomal proteins

Our analysis defined a total of 22 out of 53 mitoribosomal proteins shared with bacterial ribosomes encoded exclusively in nuclear genomes (Fig. 2, Supplementary Table S2). However, one such gene, bS6m, was recently identified in the mitochondrial genome of *Mantamonas sphyraenae* (CRuMs) (69). This is surprising, as all other eukaryotes, including another member of CRuMs, encode bS6m in the nuclear genome or have lost it completely. Because bS6m of *M. sphyraenae* branches with bS6m genes from other eukaryotes (Supplementary Fig. S7), we ruled out its non-mitochondrial origin. Similar to bS6m, proteins uL23m and bL36m are only encoded in mitochondrial genomes of a single group, CRuMs and Malawimonadida, respectively (Supplementary Table S2). Altogether, at least 32 mitoribosomal proteins were encoded in the mitochondrial genome in LECA.

The most plausible scenario explaining the absence of the three genes from mitochondrial genomes of most eukaryotes involves independent transfers to the nuclear genome. In general, intracellular transfer of genes encoding ribosomal proteins is an ongoing and relatively common process (70) especially pronounced in plants (71), as demonstrated by several independent transfers of *uL2m* and *uL19m* genes within the group of Fabaceae (72). In our data this is further supported by the identification of at least four independent transfers of uS11m during the evolution of Archaeplastida (Supplementary Table S2).

The case of bL36m is more complex. Bacterial and plastidial bL36 proteins occur in two distinct types, referred to as p-type and c-type (73), bL36A and bL36B (74) or C- and C+ (75), which are distinguished by a c-type specific four amino-acid insertion (Supplementary Fig. S8A). Some bacteria, including *Escherichia coli*, encode both types, contributing to ribosome heterogeneity and function specialization (76). Mitochondrial bL36m proteins are almost exclusively encoded in the nuclear genome and belong to the p-type. The only exception we identified is the group Malawimonadida, which possess a c-type bL36m with phylogenetic affinity to α-proteobacterial counterparts, which is encoded in mitogenome (Supplementary Fig. S8B). Since α-proteobacteria typically do not encode p-type bL36m, the most plausible hypothesis is that ancestral mitochondrially-encoded c-type bL36m protein have been replaced by nucleus-encoded p-type bL36 in mitoribosomes of other eukaryotes, possibly acquired by multiple horizontal gene transfers. An alternative and less probable scenario is that the α-proteobacteria-like ancestor of mitochondria might have encoded both p- and c-type bL36. This situation markedly resemble fate of multisubunit mitochondrial RNA polymerase of bacterial origin that is still encoded in mitogenomes of jakobids and was replaced by single-subunit polymerase of viral origin in all other eukaryotes (77).

### Intercompartmental transfer of ribosomal proteins

Given the high conservation of ribosomal proteins of bacterial origin, nucleus-encoded ribosomal proteins retargeted to a different compartment could plausibly replace the original ribosomal component with little to no modification. Indeed, here we show that proteins bL19, bL21 and bS16 identified in cryoEM (78–80) structures of plant plastid ribosomes are paralogs of nuclear encoded mitoribosomal proteins bL19m, bL21m, and bS16m. The genes encoding original chloroplastic bL19c and bL21c have been lost, and the respective proteins were replaced by the counterparts of mitochondrial origin (Fig. 4). The gene encoding bS16c is retained in some organisms, including *A. thaliana*, but the protein itself is not present in any resolved structures. In other plants, the gene has been lost completely, indicating that *A. thaliana* represents an intermediate state (81) (Table 2). Our data further indicate that the replacement occurred at different points of evolution as genes *bL19m* (Supplementary Fig. S9) and *bL21m* (Supplementary Fig. S10) were duplicated in the common ancestor of embryophytes and angiosperms, while the duplication of *bS16m* occurred independently in the lineages of *Setaphyta* and eudicots (Supplementary Fig. S11). Finally, the ancestor of the lycophyte *Selaginella* most likely utilized a paralog of uL33m, predicted to possess a plastid-targeting signal, instead of uL33c, which has been lost in all three species we investigated (Supplementary Fig. S12).

**Figure 4.**
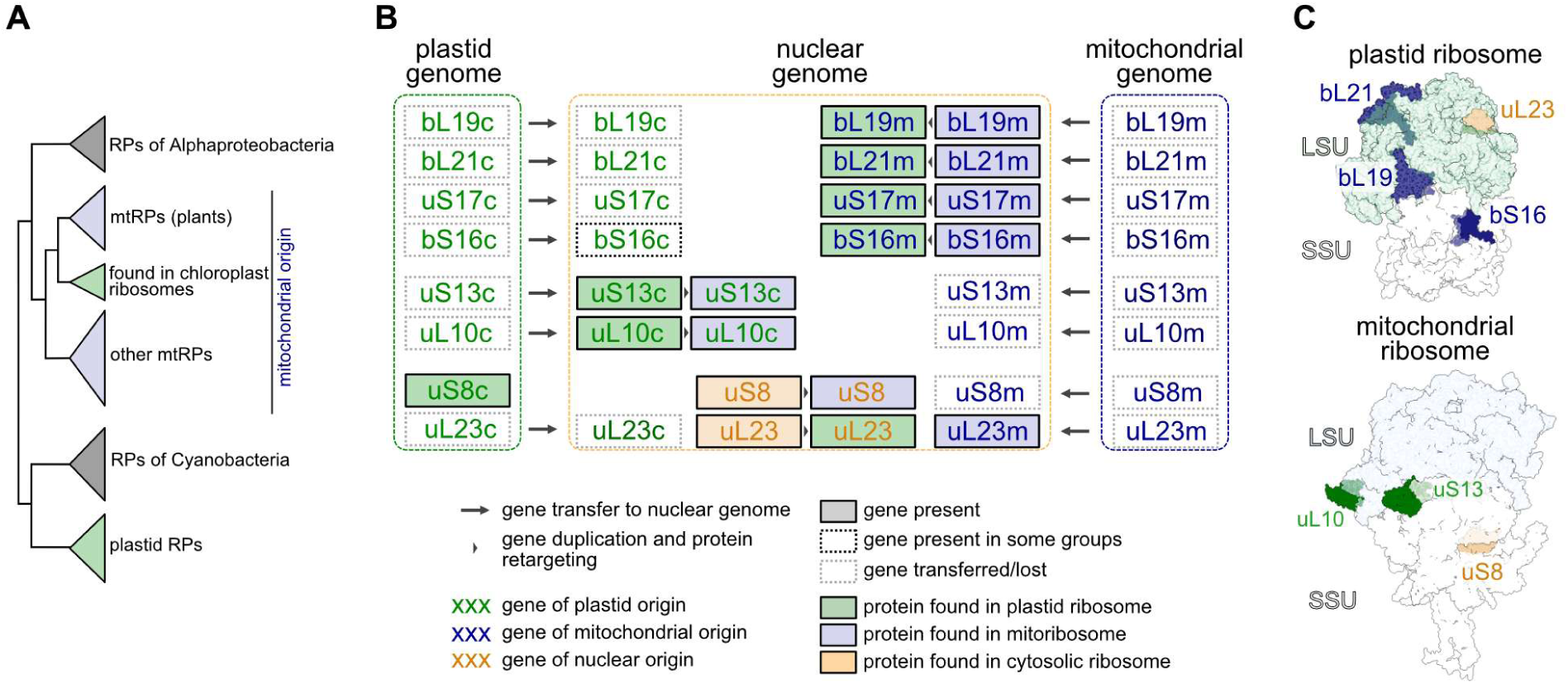
Several proteins of organellar ribosomes in plants were replaced by homologs originating in a genome from a different cellular compartment. **(A)** A schematic phylogenetic tree documenting mitochondrial origin of a protein found in the plastid ribosome of plants. **(B)** The evolutionary history of genes encoding selected ribosomal proteins in seed plants or their subgroups. Gene transfers, duplication events, re-targeting and losses are indicated. **(C)** Structures of plant organellar ribosomes with proteins originating in a genome of other cellular compartments highlighted.

**Table 2.** List of proteins transferred between different types of ribosomes.

| Replacement of original protein by retargeted protein with different origin |  |  |  |  |
| --- | --- | --- | --- | --- |
| Protein | Present in | Origin | Organisms | References |
| uL10 | mitoribosome | plastidial | Brassicales, Caryophyllales, Monocots | (94,95), this study |
| bL19 | plastid ribosome | mitochondrial | Embryophyta | this study |
| bL21 | plastid ribosome | mitochondrial | Angiosperms, *Mesostigma* * | (96,97), this study |
| uL23 | plastid ribosome | cytosolic | Caryophyllales, Cuscutaceae | (79,85), this study |
| uL24 | plastid ribosome | cytosolic | Euglenida, Eustigmatophyta | (86,87) |
| uL33 | plastid ribosome | mitochondrial | *Selaginella* | this study |
| uS8 | mitoribosome | cytosolic | Tracheophyta | (83,84), this study |
| uS13 | mitoribosome | plastidial | Rosids, Santalales | (83), this study |
| uS15* | plastid ribosome | mitochondrial | *Anthoceros*, *Selaginella* | this study |
| bS16 | plastid ribosome | mitochondrial | Setaphyta, Cuscutaceae, *Populus*, *Medicago* | (81,88,89), this study |
| uS17* | plastid ribosome | mitochondrial | Viridiplantae | this study |
| uL2, uL14, uS12* | plastid ribosome# | mitochondrial | Apicomplexa | (82) |
| Co-existence of original and retargeted gene and protein |  |  |  |  |
| Protein | Present in | Origin | Organisms | References |
| uL23 | plastid | cytosolic | Euasterids | this study |
| bS16 | plastid | mitochondrial | *Arabidopsis*, *Ricinus*, *Glycine* ** | (81,88,89), this study |

\*tentative retargeting
\*\*duplication event in Eudicots with subsequent gene loss among many lineages
#encoded in plastid genome

We further identify two additional cases of possible inter-compartmental replacements. Specifically, uS17 identified in the plastid ribosomes in *A. thaliana* and its orthologs from Chloroplastida and lineages with Chloroplastida-derived plastids, branch together with uS17m (Supplementary Fig. S13). Similarly, the predicted targeting of one nuclear copy of *uS15m* to the plastid, paired with the absence of uS15c in hornworts and lycophytes strongly suggests its retargeting (Supplementary Fig. S14). However, the relevant uS17 and uS15 sequences exhibit long branches in the phylogenetic tree, and statistical support across the tree is generally low. Therefore, we categorize these two replacements as tentative, together with the cases of *uL2*, *uL14* and *uS12* in plastid genomes of Apicomplexa, previously proposed to be transferred from mitochondrial genomes (82).

Our analyses also support several reported cases of inter-compartmental retargeting. A paralog of cytosolic uS8 replaced the original uS8m in mitochondrial ribosomes of Tracheophyta (83,84) (Supplementary Fig. S15) and paralogs of plastidial uS13c (Supplementary Fig. S16) and uL10c (Supplementary Fig. S17) substituted their mitochondrial counterparts in *Arabidopsis*(*71*), potentially three times independently. Similarly, a paralog of cytosolic uL23 (Supplementary Fig. S18) replaced its plastic counterpart in some flowering plants (85). Further, a paralog of cytosolic uL24 has been engaged in plastidial ribosomes independently in photosynthetic euglenids and Eustigmatophyta (86,87).

In two cases, the original and re-targeted proteins co-occur in the same compartment, likely representing an intermediate state before the original component is fully replaced. This is based on *in silico* prediction of protein localization (paralog of cytosolic uL23 in plastid in euasterids, Supplementary Fig. S18) or by experimental validation (one of the bS16m proteins of mitochondrial origin serves in the plastid in several lineages of seed plants; Supplementary Fig. S11 and references (81,88,89)). A list of all replacements that were previously published or revealed in this study is provided in Table 2.

Altogether, our results document compositional crosstalk between ribosomes from all compartments (Fig. 4, Table 2). This phenomenon may be more widespread, but individual cases can be confirmed only through experimental approaches. The ability of ribosomes to incorporate proteins of evolutionarily distant origin underscores their structural conservation across the tree of life, despite substantial sequence divergence. In this context, it is tempting to speculate that some proteins have dual localization, participating in ribosomes in two different cellular compartments but to our knowledge, no such cases have been reported to date.

## Conclusions

Our results provide comprehensive insight into the distribution of mitoribosomal proteins across eukaryotes, and into the evolutionary trends that shaped mitochondrial ribosomes. The period between the endosymbiotic event and the emergence of LECA was characterized by a net protein gain despite minor to no changes in the rRNA composition. After the radiation of eukaryotes into extant supergroups, many proteins (including those of bacterial origin) were lost in specific lineages, in most cases multiple times independently. Nevertheless, the fact that all structurally characterized mitoribosomes contain at least two proteins unique to that structure (Fig. 2) indicates that the trend in acquisition of novel proteins is ongoing in most lineages, albeit at varying rates. Similarly, while reduction of ancestral mitochondrial rRNA is ongoing and convergent in nearly all groups, it differs in extent. In addition, rRNA was expanded locally by accretion in some cases (9,14), and some structures show both minor reductions and accretions (8,16). Consequently, mitoribosomes of certain groups are highly conserved in terms of RNA architecture and protein content, whereas others are derived. The former group includes vascular plants and jakobids (this study and (20)), and the latter contains euglenozoans, alveolates, and opisthokonts. Clade-specific mitoribosomal proteins often contain folds of enzymes or proteins abundant in the mitochondrial matrix, documenting that while entrenchment contributes substantially to the evolution of mitoribosomes, it is mostly unguided and proliferates primarily through neutral means. This is in line with the view that the general diversification of mitoribosomes may be a result of constructive neutral evolution (90,91). As an inherent limitation of our *in silico* approach, we are unable to detect ribosomal proteins specific for certain groups or organisms or presence of proteins localized in two types in ribosomes in the same organism; such proteins can be identified only experimentally.

Our study is based on thorough analysis of 121 organisms, 85 of them represented by high-quality data from organellar as well as nuclear genes. Considering the differences observed even among relatively closely related species, illustrated by the presence of uS7m in *A. thaliana* and its absence in both *Nicotiana* species, deeper sampling would likely reveal even greater diversity and further support the high evolutionary plasticity of mitochondrial ribosomes in extant lineages. It remains to be explored whether the same is true also for plastid ribosomes. A recent structural study has already highlighted significant architectural variation (92), indicating together with the shared evolutionary environments of ribosomes from the two organelles, that there still exists a wealth of diversity to be discovered.

## DATA AVAILABILITY

All data from this study are available as Supplementary materials or have been deposited to Figshare under accession number XXX.

## SUPPLEMENTARY DATA

Supplementary data are available at *NAR* Online.

## AUTHOR CONTRIBUTIONS

Conceptualization: T.P., V.T., O.G. Methodology: T.P., J.Ř., V.T., O.G. Investigation: T.P., J. Ř., Pi.C., E.Z., J.E.W., V.T., O.G. Data Curation: T.P., J.Ř., Pi.C., V.H., A.S., Pr.C., J.E.W., A.S.P., K.S., V.T. Visualization: T.P., J.Ř., Pr.C., K.S., V.T., O.G. Software: V.H., A.S., A.S.P. Funding acquisition: T.P., A.S.P., V.T., O.G. Supervision: T.P., A.S.P., V.T., O.G. Writing - original draft: T.P., V.T., O.G. Writing - review & editing: All authors.

